# Modelling dynamics of human NDPK hexamer structure, stability and interactions

**DOI:** 10.1101/2024.09.19.613900

**Authors:** Yee Ying Lim, Kedar Nath Natarajan

## Abstract

Nucleoside diphosphate kinases (NDPKs) are evolutionarily conserved multifunctional enzymes involved in energy metabolism and gene regulation. NDPKs primarily regulate nucleotide pool turnover by catalyzing the transfer between nucleoside triphosphate (NTPs) and their deoxy derivatives, maintaining cellular homeostasis. The NDPK hexameric assembly is needed for kinase activity, but its precise assembly into homo-/hetero-oligomeric complexes remains poorly understood. How quaternary structure affects NDPK activity is limited by high subunit homology, experimental challenges in isolating *in vivo* heterohexamers and subunit abundances across cellular compartments. We identify conserved Arg27 across group I NDPKs (NME1-4) as the key residue for hexamer assembly. The Arg27 ensures similar hexameric assembly across subunits and mediates inter- and intra-molecular monomeric interactions, while Arg27 mutation leads to decreased binding affinity, dynamics, and complex destabilization. The double and triple Arg NME4 mutations destabilize hexamer into dimer due to shorter C-terminal region. Simulating NME1-3 with Arg mutations and shortened C-terminal recapitulates hexameric destabilization, highlighting role of the C-terminal region in stabilizing NDPK hexamers. Comparing heterohexameric complexes, we report NME1-NME2 (A_1_B_5_) complex as most stable and abundant, owing to predominant subunit nuclear localization. We propose that Arginine residues, C-terminal sequence and subunit abundances contribute to formation and stabilization of NDPK heterohexameric complexes.

## Introduction

Nucleoside diphosphate kinases (NDPKs) are evolutionarily conserved, ubiquitous multifunctional histidine kinases that play a crucial role in nucleotide metabolism and diverse cellular processes [1]. Their nomenclature is based on enzymatic function, *i.e.*, NDPK activity, but they are also referred to as non-metastatic enzymes (NME/NM23), due to their role in metastasis suppression across multiple tumour types [2–5]. NPDKs catalyse the transfer of high-energy ɣ-phosphate from a nucleoside triphosphate (NTP) to a nucleoside diphosphate (NDP) through a ping-pong mechanism and are critical components of cellular metabolism, maintaining cellular homeostasis [6]. Ten NME subunits are found in humans (NME1–10), categorized into group I and II NDPKs based on their enzymatic activity, subcellular localization, and evolutionary similarity. The group I NDPKs (NME1-4) monomers are small proteins (15-18 kDa) with high sequence similarity, contain a single NDPK catalytic domain, and vary in length of N-terminus and C-terminal sequences (∼10-15 amino acids) [7]. The group II members consist of divergent sequences with either no or low NDPK activity [8]. The group I NDPKs exhibit conserved NDPK activity, functioning through conserved histidine residue, and near identical active sites [9–11]. The NME4 is unique as it contains an N-terminal mitochondrial localization sequence and a shorter C-terminal region [12]. Overall, eukaryotic NDPKs are crucial for a variety of cellular processes, including epithelial-mesenchymal transition [13], G-protein signal transduction [14], plasma membrane remodelling [15, 16], regulation of gene expression [17] and Wnt [18], TGF-β [13], MAPK/ERK signalling [19].

Eukaryotic NDPKs are predicted to function as monomers [20, 21], dimers [22], trimers [23], hexamers [24], with their oligomeric forms important for diverse biological functions. The purified homohexameric crystal structures reveal a shared D3 symmetry with conserved histidine residue (NME1/2 H118; NME3 H135; NME4 H151) in the cleft, governed by a “clamp” formed between the *Kpn* loop and helical αA-α2 hairpin loop (Fig. S1A) [25–29]. Several oligomers can co-exist in response to dynamic cellular microenvironment; for example, elevated oxidative stress promotes NME1 dimerization via disulfide bond formation between C4 and C145 [30], while NME1 mutations (P96S or S120G) destabilise hexamer to dimer due to disturbed interaction between *Kpn* loop and the C-terminal of the adjacent monomer, coupled with loss of NDPK [31, 32]. Of the group II NDPKs, NME6 is the only ubiquitous monomeric subunit, but lacks NPDK activity [20, 33, 34].

The highly abundant NME1 and NME2 subunits have 88% sequence similarity, can catalyse NDPK functions, and are localised both in the cytoplasm and nucleus [35]. The distinct ∼18 residues contribute to different monomeric isoelectric point (pI), resulting in an acidic NME1 and basic NME2 [36–38]. Their distinct relative abundance across cellular compartments and affinity likely contribute to diverse cellular roles. For instance, NME1 is a stronger metastasis suppressor (than NME2) in cancer cells, has a lower affinity to double-stranded DNA, exhibits 3’-5’ exonuclease activity, and contributes to insulin secretion and endocytosis [39]. On the other hand, NME2 binds with folded DNA structures and regulates promoter accessibility and transcription in several cancers and stem cells [21, 40–42]. Surprisingly, both NME1 (group I) and NME7 (group II) are reported to function as extracellular growth factors through cell-surface receptors in stem cells, despite high sequence diversity [43, 44]. It remains unclear how much sequence diversity (pI) and oligomeric states contribute to different biological roles. Since the ∼18 differing residues between NME1 and NME2 are located outside of the hexamer interacting sites, they result in indistinguishable heterooligomeric complexes and intra- molecular interactions [24]. A recent work used native mass spectrometry and spectral modelling to suggest heterohexameric NME1-2 nuclear states, despite both subunits lacking nuclear localization signals [24]. Thus far, various oligomerization states have been reported for group I NDPKs, including homohexameric crystal structures from *in vitro* overexpression constructs, but the link between oligomerization, complex stability, abundance, and broad intracellular NME function has been elusive. No heterohexameric crystal structures have been purified due to experimental challenges with high sequence similarity, lack of specific antibodies, and analytical approaches to study *in vivo* acid-labile protein histidine phosphorylation and indistinguishable homohexameric forms [22, 24, 42, 45, 46]. While *in vitro* over-expression coupled with native mass spectrometry and PAGE, has identified homohexamers, these do not reflect *in vivo* dynamic configurations [24]. Therefore, we propose *in silico* structural modelling of experimental data, perturbation, and careful interpretation towards an improved understanding of group I NDPK states and biological function. Here, we employ structural priors with alanine scanning mutagenesis (CASM) to identify key residues in all group I NDPK homohexameric assemblies and report the contribution and stability of individual monomers towards forming homo- and heterohexamer complexes. We select parameters and simulations to mimic cellular physiological conditions and capture time-dependent protein interactions, which are difficult to extrapolate through *in vivo* crystal structure analysis. By modelling possible heterohexameric configurations and integrating molecular dynamics (MD) simulations, we reveal the NME1-NME2 (A_1_B_5_) complex as likely nuclear heterohexamer state.

## Results

### Homohexameric assembly of NDPK group I subunits

To assess the homohexameric assembly of group I NDPK proteins, we first compared the monomers and their tertiary folding. All group I NDPKs consist of four strands of stacked antiparallel *β* sheets (*β_2_β_3_β_1_β_4_*) to form a central core, surrounded by eight *α* helices, with each subsequent *β* sheet followed by two *α* helices (Fig. S1A). Comparing the atomic distances between superimposed group I monomers shows a similar folding with resultant backbone C_α_ root mean square deviation (RMSD) <0.5Å, with the strongest overlap between NME1 and NME2 (RMSD = 0.23Å, Fig. S1B). Of note is NME4, which harbours a 33 amino-acid mitochondrial localization signal (MLS) in its N-terminal region, which is cleaved upon reaching the mitochondrial matrix. Consistently, the RMSD difference between NME4 and NME1 is slightly larger (RMSD = 0.46Å), but folding is quite similar between all group I NDPK isomers.

We employed available crystal structure coordinates to compare oligomeric group I complexes, arguing against reconstruction of sequence-based monomers due to suboptimal solutions. Despite the differences in length of N- and C-terminal sequences length across group I monomers, their available crystal structures present a highly similar homohexameric quaternary arrangement with a D3 symmetry. This int turn can be visualized as a dimeric layer of stacked trimers (Fig. S2A-B, Supplementary Table 1). Between the bilayer trimers (‘A-B-C’, ‘D-E-F’, Fig. S2B), the dimers interact with the largest burial surface area, involving two layers of antiparallel *β* sheets interacting with each other [47]. Meanwhile, each of the bilayer trimers is assembled to have three monomers, with three possible dimer pairs (‘A-B’, ‘A-C’, ‘A-D’, Fig. S2C), arranged in a head-to-tail orientation, and ∼6-17 C-terminal residues forming intermolecular interactions with the *Kpn* loop (residue 94- 114) of the adjacent monomer (Fig. S2D). This results in a protein burial surface area of 300Å^2^, which represents ∼30% of the total surface area of a monomer, indicating that the C-terminal region could contribute towards stabilizing the homohexameric complex at the trimer interface (Fig. S2D).

We next assessed which residues facilitate interaction between monomers and confer flexibility and stability to the homohexamers. We performed a residual interaction map for each pair of interacting monomers (‘A-B’, ‘A-C’, ‘A-D’, Fig. S2C), which becomes repeated in the stacked trimer layer. The NME1-4 homohexamers show a short range of C_α_ RMSD (∼0.9-1.6Å), indicating highly similar quaternary structural assembly and dynamics (Fig. 1A). Superimposition of NME2, NME3, and NME4 onto the NME1 homohexamer reveals a near identical assembly with an RMSD deviation of 0.96Å (NME2), 1.12Å (NME3), and 1.64Å (NME4) (Fig. 1B). To assess the stability of the homohexameric complex in a solvated system, we calculated and compared the total energy of each complex. We find NME3 homohexamer is the most stable construct (TE: -1.62×10^5^ kcal/mol), followed by NME2 (TE: -1.57×10^5^ kcal/mol), NME4 (TE: -1.54×10^5^ kcal/mol), and NME1 as least stable (TE: -1.47×10^5^ kcal/mol) (Fig. 1B). Despite higher sequence similarity between NME1 and NME2, we find that the difference in total energy between NME3 and NME1 is as high as 14,863 kcal/mol. NME1 and 3 carry similar kinetic energy (∼1.9 x 10^5^ kcal/mol); however, conserved covalent bonds in NME3 contribute to its lower potential energy and higher stability (than NME1).

**Fig. 1.**
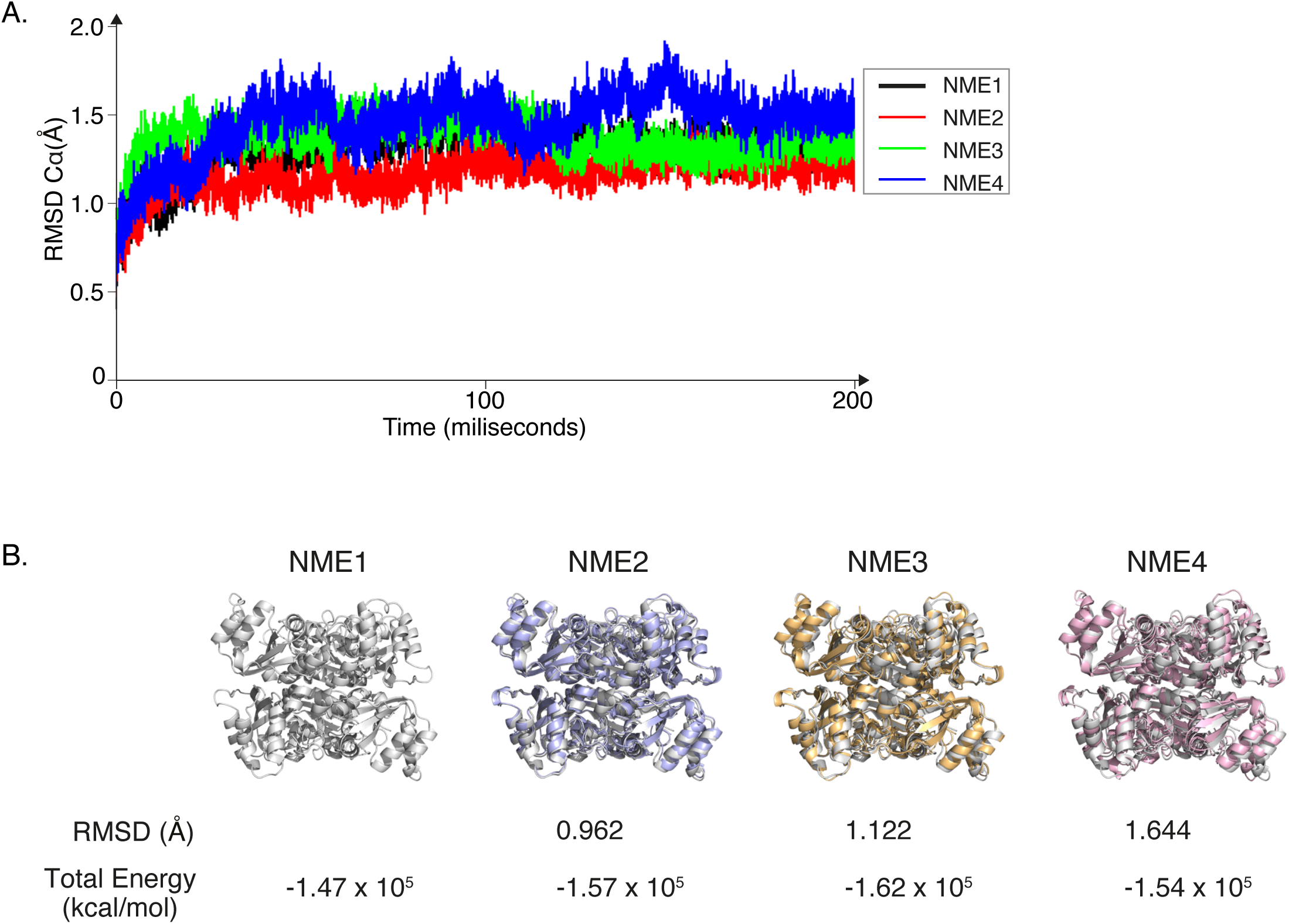
MD simulation of group I NPDK homohexamers. **(A)** The backbone C_α_ RMSD of each NME hexamer against time, with NME1 (black), NME2 (red), NME3 (green) and NME4 (blue).B Hexamer superimposition of other NDPKs (NME2: blue, NME3: orange, NME4: pink) on NME1 (grey), with the respective backbone C_α_ RMSD and total energy (kcal/mol) is listed underneath the superimposed structures.

To capture which residue contributes to structural flexibility and interaction with external molecules (e.g., ligands or DNA), we performed root mean square fluctuation (RMSF) analysis. We find NME1 residues 54- 57 and 94-98 (*α*_A_-*α*_2_ hairpin and *α*_3_-*β*_4_ *Kpn* loop) showed increased flexibility in forming a “clamp” that enables external ligand accessibility (e.g., Acetyl CoA) into the *β_4_* sheet binding cleft (Fig. S2D). Conversely, NME2, NME3, and NME4 homohexamers show lesser *Kpn* region flexibility, which indicates a restrictive access to the active site. The structural rigidity is particularly critical for NME2, as it exposes positively charged residues at the surface to enhance DNA recognition and binding (as opposed to NDPK activity). Furthermore, to examine homohexameric stability, we calculated binding free energy (MMGBSA) for all group I NDPKs and found that monomer binding affinity is consistent with the hexamers’ total energy. We report that the NME3 hexamer complex is the most stable among group I NDPKs (ΔΔG_bind_ = -267.91.91 kcal/mol, Table 1), followed by NME2 (ΔΔG_bind_ = -149.63 kcal/mol), NME4 (ΔΔG_bind_ = -148.26 kcal/mol), and NME1 (ΔΔG_bind_ = -147.43 kcal/mol, Table 1). As NME4 uniquely consists of a truncated C-terminal region, we repeated the MMGBSA calculations on truncated NME1, NME2, and NME3 (Δ10ct). All truncated homohexamers show an increased binding energy relative to their wildtypes, with the most prominent increase in NME3 Δ10ct (ΔΔG_bind_ = 178.52 kcal/mol), followed by NME2 Δ10ct (ΔΔG_bind_ = 71.10 kcal/mol) and NME1 Δ10ct (ΔΔG_bind_ = 40.73 kcal/mol, Table 1). The large differences in binding energy in truncated variants (NME1 Δ10ct, NME2 Δ10ct, and NME3 Δ10ct) indicate C-terminal regions contribute significantly to homohexamer assembly, particularly at the trimer interface. Surprisingly, the increased binding energy of truncated variants did not result in destabilized homohexamers (Table 1), suggesting that the C-terminal region is regulated by additional determinants (melting point, pH) in mediating the homohexameric complex.

**Table 1.**
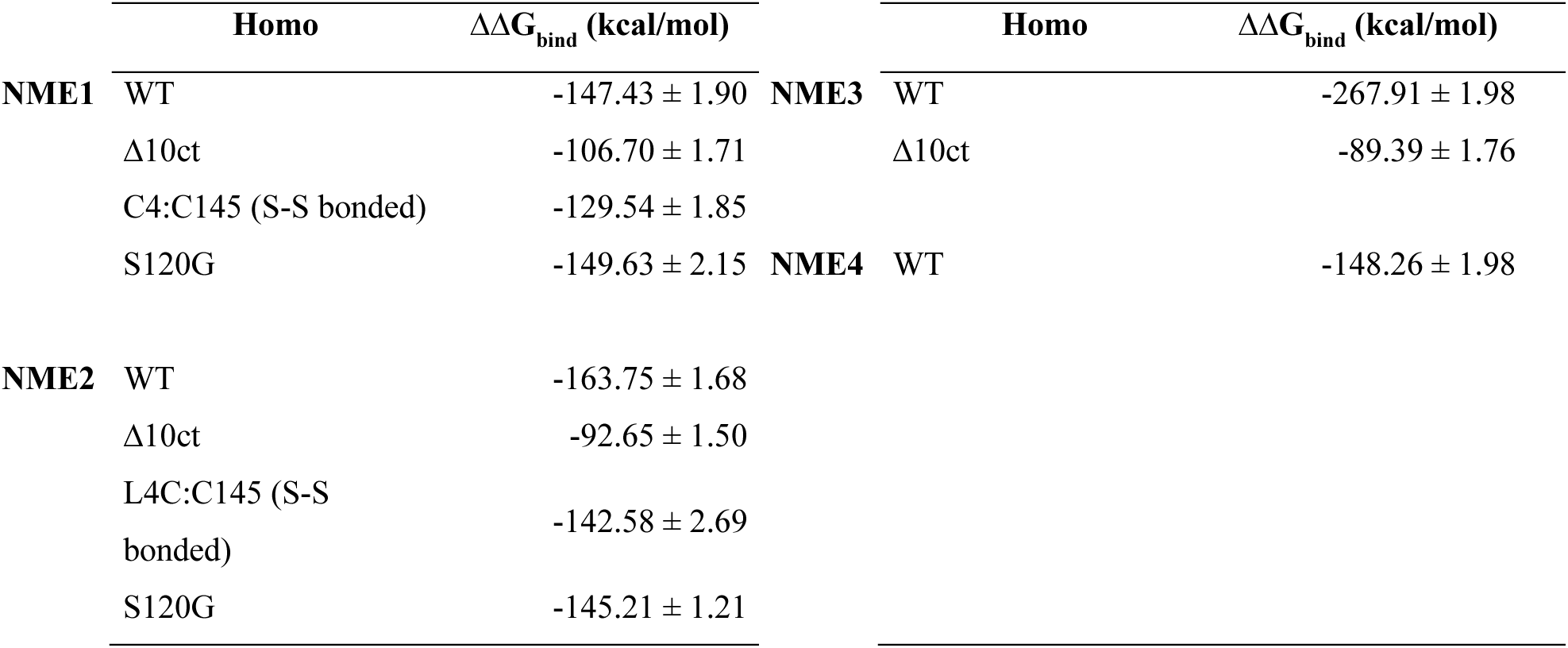
MMGBSA calculations for binding free energy changes in NME homohexamers.

| | Homo | $\Delta\Delta G_{\text{bind}}$ (kcal/mol) | | Homo | $\Delta\Delta G_{\text{bind}}$ (kcal/mol) |
| --- | --- | --- | --- | --- | --- |
| **NME1** | WT | $-147.43 \pm 1.90$ | **NME3** | WT | $-267.91 \pm 1.98$ |
| | $\Delta 10\text{ct}$ | $-106.70 \pm 1.71$ | | $\Delta 10\text{ct}$ | $-89.39 \pm 1.76$ |
| | C4:C145 (S-S bonded) | $-129.54 \pm 1.85$ | **NME4** | WT | $-148.26 \pm 1.98$ |
| | S120G | $-149.63 \pm 2.15$ | | | |
| **NME2** | WT | $-163.75 \pm 1.68$ | | | |
| | $\Delta 10\text{ct}$ | $-92.65 \pm 1.50$ | | | |
| | L4C:C145 (S-S bonded) | $-142.58 \pm 2.69$ | | | |
| | S120G | $-145.21 \pm 1.21$ | | | |

We next performed MD simulations on experimentally tested mutations in NME1 and NME2 to decipher the affinity contribution of each monomer in their homohexamer configurations. These include S120G and the C4:C145 disulphide bridge, both of which result in a dimeric complex in a high oxidative stress environment [48] . Notably, NME2 contains a Leucine at position 4, which we mutated to Cysteine to mimic the constrained disulphide bond. Across simulations, the S120G mutation did not drastically alter the binding energy of NME1 (ΔΔG_bind_ = -149.63 kcal/mol) or NME2 (ΔΔG_bind_ = -145.4213 kcal/mol, Table 1). However, the constrained disulphide bond reduced the stability of the complex with an increased binding energy (∼20 kcal/mol) for both NME1 and NME2 (Table 1). Our observations suggest a critical threshold (∼-140 kcal/mol) for monomeric contribution, beyond which homohexameric assembly is destabilised.

### Arginine residue is critical for homohexamer interactions and assembly

We next set out to identify the key residues for monomeric intermolecular interactions within the homohexamers, using an alternative approach. Through alanine scanning mutagenesis, we scanned and mutated all NME1-4 residues to Alanine and computed shifts in binding free energy at all protein interfaces (‘A-B’, ‘A-C’, ‘A-D’, Fig. S3A-D). Notably, we repeat the mutagenesis by retaining residues that lack altered binding energy so as to preserve the proximal distances between neighbouring monomers. We first generated interactions maps (from PDBsum) to report on monomeric contribution to the amount and position of residues in the wildtype hexameric assembly (Fig. S4A-B, S5A-B). After performing unconstrained molecular dynamics (MD) and mutagenesis, we regenerated a residue interaction map for all group I NDPK isomers and used the backbone C_α_ average RMSD to evaluate structural deviations across time (Fig. 2, Fig. S6A-B, S7A- B). The mutagenesis predicts positionally conserved Arginine residue (NME1: Arg27, NME2: Arg27, NME3: Arg44, NME4: Arg60) to be important across group I NDPKs. The Arginine residues have adhesive characteristics that can mediate strong intermolecular interaction [49, 50]. Notably, the conserved Arginine residue resides in a manner that it has close proximity across all interfaces, influencing the integrity stability of hexameric assembly (Fig. 2, Fig. S3A-D). Multiple sequence alignment between group I and group II NDPKs reveal that NME5 and NME6 lack arginine residue and consequently cannot form hexameric complexes. We next performed site-saturated mutagenesis on group I NDPKs; capturing Arg27Pro mutations causes maximum destabilizing, while Arg27Ala mutation has least impact on homohexamer assembly. We performed a systematic single-residue permutation on all group I NDPKs (Arg27 NME1, Arg27 NME2, Arg44 NME3, Arg60 NME4) to both Proline and Alanine, followed by MD simulation (Fig. 3A-B, 4A-B). Across the MD simulations, single mutations (Arg to Pro; Arg to Ala) did not disrupt homohexamer configuration but reduced the compactness of the homohexamers, as highlighted by increased total energy and C_α_ RMSD with expanded configuration from the hexamer core (compared to wildtype, Fig. 3B). The Arg-to-Pro substitution exposed the monomer distance away from the hexamer core while contributing to some interaction for the hexamer conformation, leading to diminished stability compared to the wildtype (Fig. 3B, Fig. S8A-B).

**Fig. 2.**
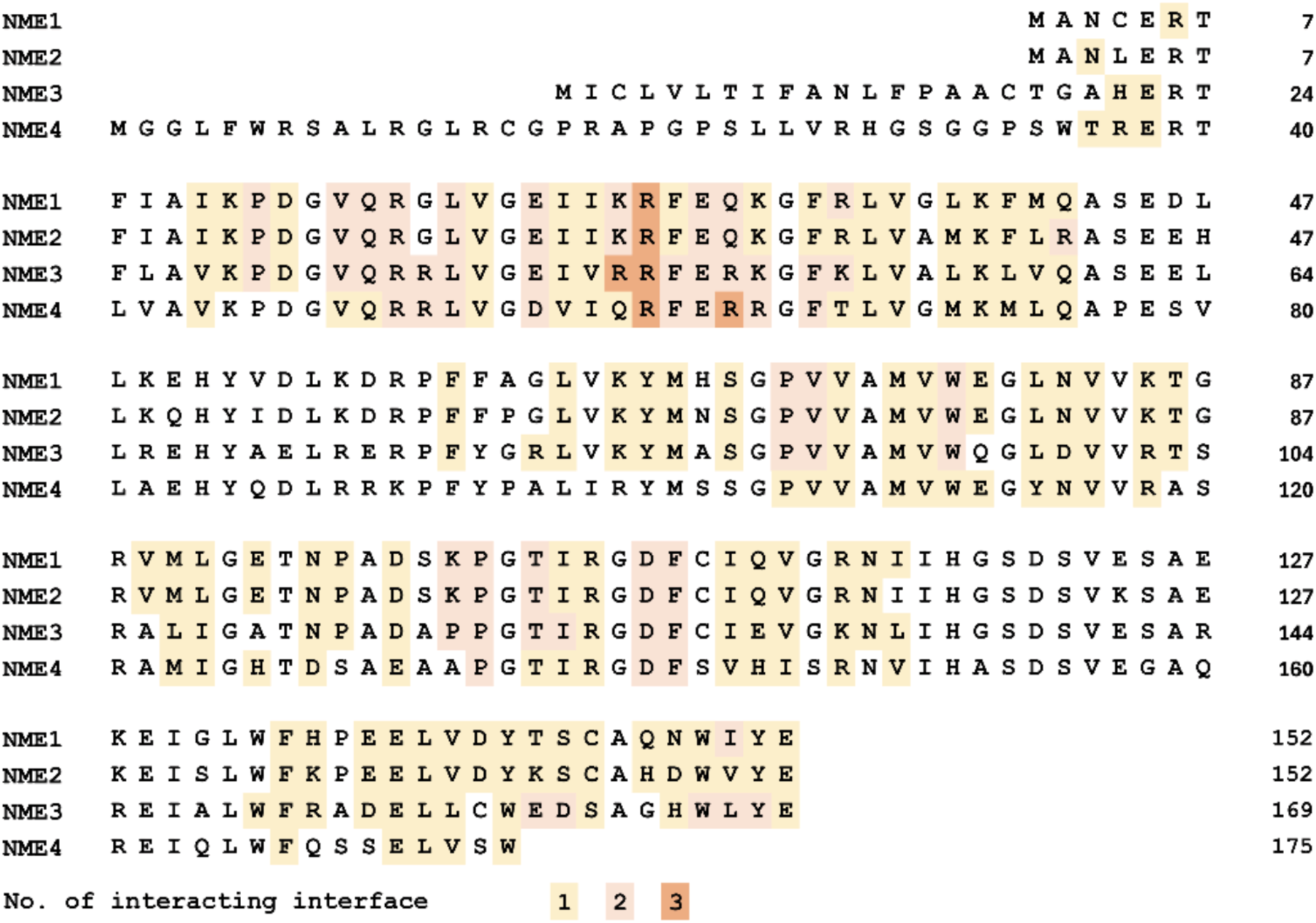
Conserved and critical residues across group I NDPK homohexamers. **(A)** CASM NME hexamers reveal key residues conserved and critical across group I NDPKs. The color coding indicates contribution of residues in mediating interaction between different interfaces (AB, AC, AD); with orange indicating 3, Pink indicating 2 and Yellow indicates 1 interacting interface. The orange regions are most critical for overall stability of hexamers as they participate in all three types of dimer interfaces.

**Fig. 3.**
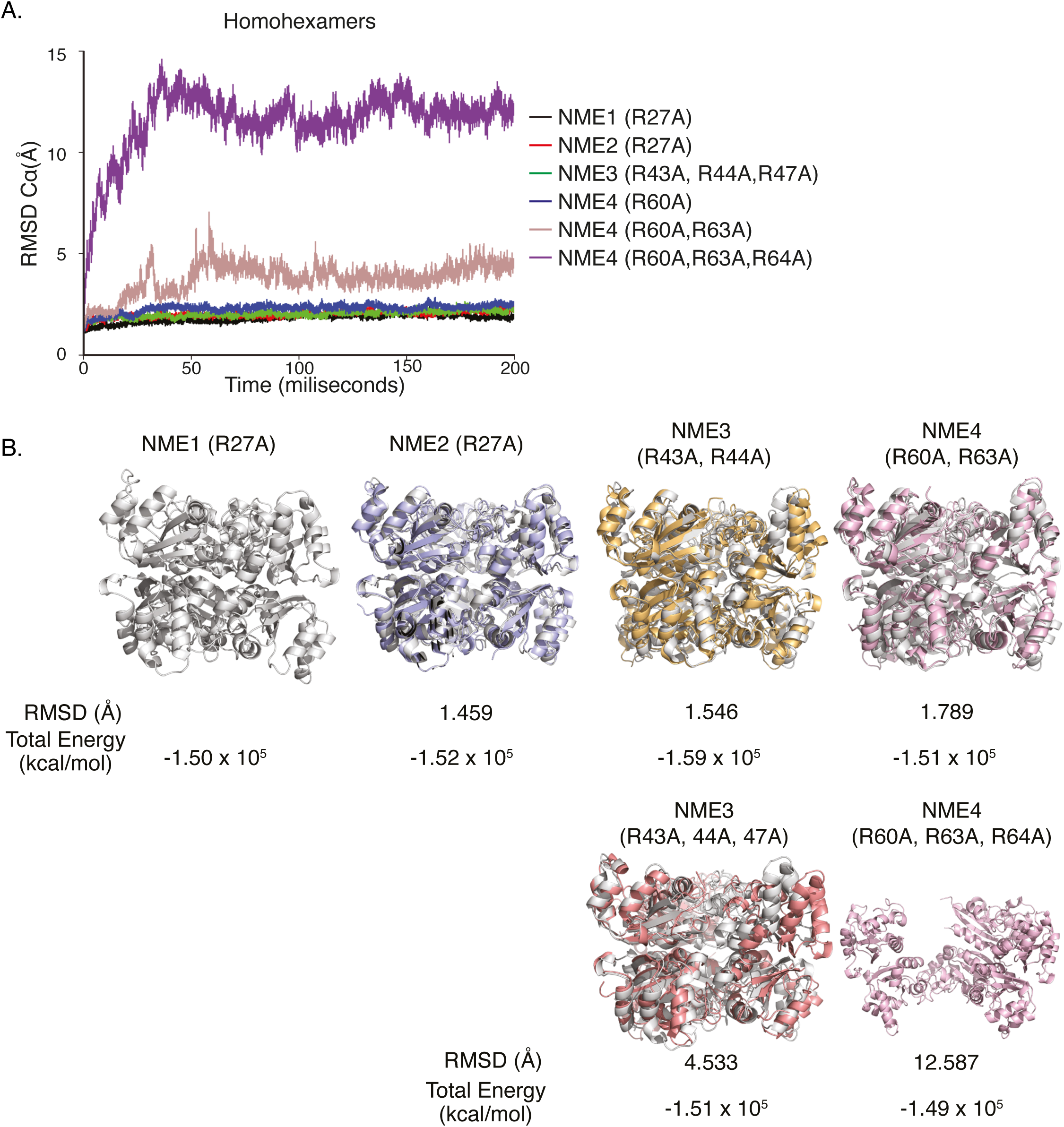
Critical Arginine mutation to Alanine in group I NDPKs affects NME4 homohexamer assembly. **(A)** The backbone C_α_ RMSD of each Arg to Ala mutated NME hexamer against time, with NME1 (Arg27Ala, black), NME2 (Arg27Ala, red), NME3 triple mutation (Arg43Ala, Arg44Ala, Arg47Ala, green) and NME4 single (Arg60Ala, purple), double (Arg60Ala, Arg63Ala, brown) and triple mutations (Arg60Ala, Arg63Ala, Arg64Ala, violet).B Hexamer superimposition of mutated NME isomers on top of mutated NME1 (Arg24Ala), with respective backbone C_α_ RMSD. The backbone C_α_ RMSD and total energy (kcal/mol) is listed underneath the superimposed structures.

Given the lack of hexameric disassembly, we reasoned that a single Arg mutation might not be sufficient, as sequence diversity might impose higher energy requirements to break intermolecular interaction within complexes. We next assessed the contribution of neighbouring Arginine or proximal residues to homohexamer assembly. We performed double and triple mutations (within a 5-residue proximity of NME1 Arg27 to Alanine and Proline), resulting in mutations on NME3 (double: Arg43, Arg44; triple: Arg43, Arg44, Arg47) and NME4 (*double*: Arg60, Arg64; *triple*: Arg60, Arg63, Arg64). Both double and triple mutations in NME4 disrupted the hexamer into a dimer within simulated time, irrespective of mutation to Alanine or Proline (Fig. 3A-B, 4A-B, Table 2). The NME4 triple mutations (Arg60, Arg63, and Arg64 to Alanine) showed a C_α_ RMSD >7Å, while perturbation was less pronounced in double mutations (Arg60 and Arg64 to Alanine) (Fig. 3B). The triple and double NME4 mutations to Proline followed a similar trend, indicating rapid hexamer destabilization to dimer constructs in NME4 triple Arginine mutation (Fig. 4B). The structural evaluation agreed with the deconstruction of NME4 hexamer into three dimer pairs (A-D configuration). The binding free energy suggests that hexamer destabilization favouring dimer formation may occur at ΔΔG_bind_ ∼ -100 kcal/mol, whereas rapid dimerization (A-D configuration) may occur when binding energy falls to ΔΔG_bind_ ∼ -60 kcal/mol (Fig. 3B, Fig. 4B, Table 2). Notably, the NME4 triple mutation (Arg to Ala) resulting in higher binding energy (ΔΔG_bind_ = -95.78 kcal/mol) than the double mutation ((Arg to Ala, ΔΔG_bind_ = -68.89 kcal/mol) was due to the incomplete dissociation of two dimer pairs that form a tetramer (Fig. 3B, Table 2). The two dimer pairs at the trimeric interface contribute to ΔΔG_bind_ ∼ -26 kcal/mol, indicating one pair of dimer interactions at the trimer interface can be as low as ΔΔG_bind_ ∼ -13 kcal/mol. The dimer pair formation in NME4 favoured the A-D configuration over the A-B or A-C configuration, agreeing with the burial surface area of each interface.

**Fig. 4.**
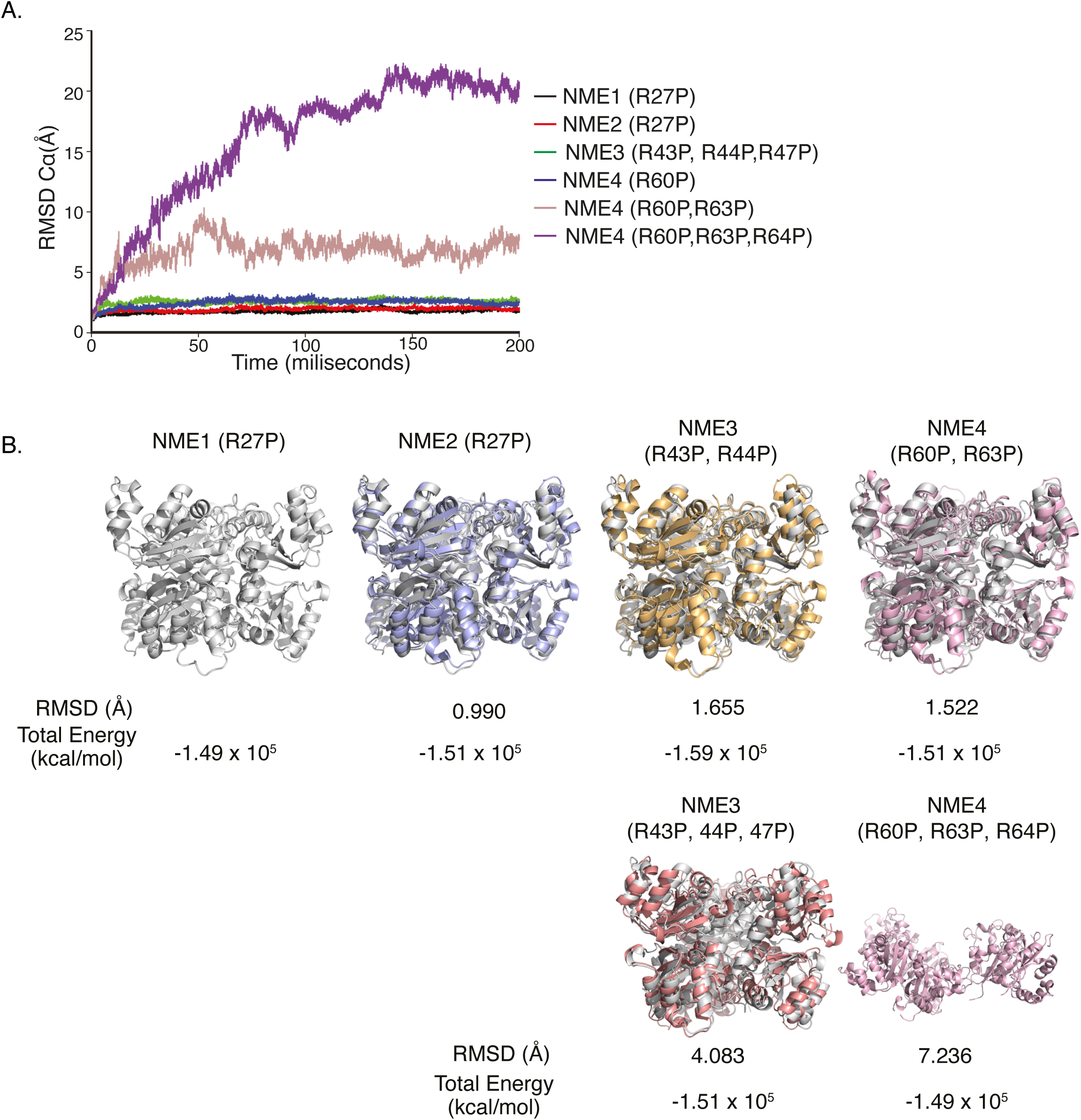
Mutation of critical Arginine to Proline residues within group I NDPK homohexamers. **(A)** The backbone C_α_ RMSD of each Arg to Pro mutated NME hexamer against time, with NME1 (Arg27Pro, black), NME2 (Arg27 Pro, red), NME3 triple mutation (Arg43Pro, Arg44Pro, Arg47Pro, green) and NME4 single (Arg60Pro, purple), double (Arg60Pro, Arg63Pro, brown) and triple mutations (Arg60Pro, Arg63Pro, Arg64Pro, violet).B Hexamer superimposition of mutated NME isomers on top of mutated NME1 (Arg24Pro), with respective backbone C_α_ RMSD. The backbone C_α_ RMSD and total energy (kcal/mol) is listed underneath the superimposed structures.

**Table 2.** MMGBSA calculations for binding free energy changes in NME mutant homohexamers.

| | Homo | $\Delta\Delta G_{\text{bind}}$ (kcal/mol) | | Homo | $\Delta\Delta G_{\text{bind}}$ (kcal/mol) |
| --- | --- | --- | --- | --- | --- |
| **NME1** | R27A | $-144.49 \pm 1.91$ | **NME3** | R44A | $-253.27 \pm 2.34$ |
| | R27P | $-155.19 \pm 2.51$ | | R44P | $-262.78 \pm 2.15$ |
| | R27A,Q30A | $-162.18 \pm 1.86$ | | R43A,R43A | $-202.04 \pm 1.72$ |
| | R27P,Q30P | $-188.86 \pm 2.11$ | | R43P,R43P | $-179.96 \pm 2.29$ |
| | R27A, $\Delta 10\text{ct}$ | $-85.19 \pm 1.21$ | | R43A,R44A,R47A | $-200.26 \pm 1.76$ |
| | Q30A, $\Delta 10\text{ct}$ | $-99.16 \pm 1.24$ | | R43P,R44P,R47P | $-163.75 \pm 1.80$ |
| | R27A,Q30A, $\Delta 10\text{ct}$ | $-85.16 \pm 1.51$ | | | |
| **NME2** | R27A | $-140.69 \pm 1.90$ | **NME4** | R60A | $-161.42 \pm 2.11$ |
| | R27P | $-156.04 \pm 1.89$ | | R60P | $-132.00 \pm 2.23$ |
| | R27A,Q30A | $-132.78 \pm 2.13$ | | R63A | $-134.76 \pm 1.77$ |
| | R27P,Q30P | $-143.32 \pm 2.12$ | | R60A,R63A | $-68.89 \pm 1.31$ |
| | R27A,Q30A, $\Delta 10\text{ct}$ | $-75.08 \pm 1.48$ | | R60P,R63P | $-131.41 \pm 2.41$ |
| | | | | R60A,R63A,R64A | $-95.78 \pm 1.63$ |
| | | | | R60P,R63P,R64P | $-67.65 \pm 1.73$ |

A rapid, significant destabilization effect in NME4 also highlights the importance of the C-terminal region by interacting with the adjacent monomer at the trimer interface. Compared to other group I NDPKs, NME4 has fewer interactions with the adjacent monomers. In contrast, NME3 with extended N- and C-terminal regions retains the highest stability, despite double or triple mutations. This stark difference between NME4 and NME3 underscores the unique importance of the C-terminal region in structural integrity. The truncated NME3 (Δ10ct) has an increased binding energy (ΔΔG_bind_ = -89.39kcal/mol), suggesting that oligomeric destabilization is amplified with a shorter C-terminal region (Table 2). Furthermore, comparing the total energy (TE) with binding free energy reveals that even the least stable NME3 homohexamers (Arg60Pro, Arg63Pro, and Arg64Pro) have lower ΔΔG_bind_ energy than the remaining wildtype group I NDPK (Fig. 4B, Table 2). Overall, our results suggest that a longer C-terminal enables overlapping Arginine interactions, which require multiple mutations at the hexameric core to disrupt the hexamer assembly.

Our results reveal a puzzling observation that C-terminal sequence diversity could impact the stability of the NPDK hexamers. The single Arginine mutations in NME1, NME2 (containing a longer C-term sequence) should decrease the overall binding energy, as predicted by site-directed mutagenesis. However, we observe that NME1 alone increases stability upon Arg27 and Glu30 mutations, in contrast to NME2 (**Table 2**). We therefore attempted to engineer NME1 and NME2 to study intermolecular affinity by imposing mutations on truncated C-terminals such that both mutants are nearly identical to NME4. The truncated C-terminal NME1 with a single mutation Arg27Ala and double mutation (R27A, Q30A, Table 2) both result in similar binding affinity (**ΔΔG_bind_** = -85.16kcal/mol), and these results are consistent for truncated NME2 with a double mutation (**ΔΔG_bind_** = -75.08kcal/mol). Notably, Arg27Ala results in higher binding energy than Glu30Ala (**ΔΔG_bind_** = -99.16 kcal/mol), supporting the mutagensis results that Arg27 is important for homohexameric assembly and stability. To study intermolecular interaction, we structurally examined nearby residues (at 5Å proximity) for their contribution to the total binding free energy in all group I NDPK. Based on distance approximation, we observe that Arg27 lies in proximity to several conserved intramolecular and intermolecular residues (Fig. S8C). Interestingly, monomer A Arg27 interacts with the monomer C Q30 in NME1 and NME2, but monomer A Arg27 interacts with monomer C R63 in NME3 and NME4. These results further highlight the importance of Arg27 in forming the trimer planar of the hexamer through interactions with Q30 in an anticlockwise manner. To assess the nature of Arg27 interactions, we repeated MMGBSA decomposition energy analysis and observed that the single (Arg27) or double mutations (Arg27, Gln30) for NME1-3 alone did not destabilize the oligomers; however, coupling mutations (single or double) with truncated C-terminals (mimicking NME4) leads to a drastic destabilization coupled with a decreased number of non-covalent interactions within homohexamers (Table 2, Supplementary Table 4-5).

### Heterohexameric assembly of NME1 and NME2

Despite the experimental challenges in purifying *in vivo* group I heterohexamers, *in vivo* acid-labile properties, and lack of distinguishing antibodies, multiple group I hetero-oligomeric complexes have been predicted to have high functional and context-dependent intracellular roles [24, 51, 52]. Here, we assess the NME1-NME2 heterohexamer configurations based on their predominant intracellular expression and localization. Despite 88% sequence similarity, NME1 and NME2 have different monomeric isoelectric point (pI) resulting in an acidic NME1 and basic NME2 (Fig. S9A). Notably, both the NDPK active site and ∼18 differing residues remain accessible and outside of the hexameric-interacting interface, such that ***(i)*** heterohexamers retain a variety of interactions and specific cellular roles as homohexamers, and ***(ii)*** the swapping of NME1 and NME2 monomers results in structurally indistinguishable and functionally distinct heterohexamers that can provide diverse interfaces for combinatorial interactions within nuclear or cytosolic compartment but also extracellular interactions (Fig. S9B). Our hypothesis is supported by the recent work, which combines theoretical modelling of native mass spectrometry data to predict heterohexamers in different cellular compartments [24].

To find the most stable configuration, we first assemble all theoretically possible combinations for NME1 and NME2, based on monomeric position and number of interfaces against monomers within a hexameric assembly (Fig. S10, Table 3, Supplementary Table 5). Of these 47 configurations, we narrowed to 11 non-redundant configurations based on the total number of interacting monomeric interfaces (A-B, A-C, A-D), resulting in pure homohexamers (A_6_B_0_, A_0_B_6_), penta-monomers (A_5_B_1_, A_1_B_5_) and three tri- and tetra-monomer configurations (A_2_B_4_, A_3_B_3_, A_4_B_2_) (Fig. S10). Repeating binding free and total energy analysis, we observe a strong preference and highest binding free energy for heterohexamer assemblies containing penta-monomer combinations (A_5_B_1_, A_1_B_5_, swapped monomer) among all tested variants (Table 3). With the exception of B_8_ (A_4_B_2_) configuration, both the affinity and binding energies of single monomers (A_5_B_1_, A_1_B_5_) are significantly stronger than native homohexamers (Fig. S10, Table 3, Supplementary Table 5).

**Table 3.** MMGBSA calculations for binding free energy changes in NME heterohexamers.

| Oligomer | $\Delta\Delta G_{\text{bind}}$<br>(kcal/mol) | Oligomer | $\Delta\Delta G_{\text{bind}}$<br>(kcal/mol) | Oligomer | $\Delta\Delta G_{\text{bind}}$<br>(kcal/mol) | Oligomer | $\Delta\Delta G_{\text{bind}}$<br>(kcal/mol) |
| --- | --- | --- | --- | --- | --- | --- | --- |
| Homohexamer |  | Tetramer |  | Trimer |  | Pentamer |  |
| **A<sub>6</sub>B<sub>0</sub>** | -147.43 ± | **A<sub>4</sub>B<sub>2</sub>** | -169.79 ± | **A<sub>3</sub>B<sub>3</sub>** | -156.92 ± | **A<sub>5</sub>B<sub>1</sub>** | -179.17 ± |
| **(NME1 WT)** | 1.90 | **(B<sub>6</sub>)** | 1.83 | **(C<sub>1</sub>)** | 2.16 | **(A<sub>1</sub>)** | 3.29 |
| **A<sub>0</sub>B<sub>6</sub>** | -163.75 ± | **A<sub>4</sub>B<sub>2</sub>** | -150.73 ± | **A<sub>3</sub>B<sub>3</sub>** | -162.80 ± | **A<sub>1</sub>B<sub>5</sub>** | -172.39 ± |
| **(NME2 WT)** | 1.68 | **(B<sub>9</sub>)** | 1.43 | **(C<sub>8</sub>)** | 1.97 | **(E<sub>1</sub>)** | 1.99 |
|  |  | **A<sub>4</sub>B<sub>2</sub>** | -169.28 ± | **A<sub>3</sub>B<sub>3</sub>** | -168.47 ± |  |  |
|  |  | **(B<sub>10</sub>)** | 2.26 | **(C<sub>10</sub>)** | 1.95 |  |  |
|  |  | **A<sub>2</sub>B<sub>4</sub>** | -149.84 ± |  |  |  |  |
|  |  | **(D<sub>6</sub>)** | 2.02 |  |  |  |  |
|  |  | **A<sub>2</sub>B<sub>4</sub>** | -142.09 ± |  |  |  |  |
|  |  | **(D<sub>9</sub>)** | 1.68 |  |  |  |  |
|  |  | **A<sub>2</sub>B<sub>4</sub>** | -159.73 ± |  |  |  |  |
|  |  | **(D<sub>10</sub>)** | 1.36 |  |  |  |  |

## Discussion

The group I NDPKs are conserved, multifunctional enzymes primarily involved in nucleotide metabolism; the oligomers function as phosphohistidine kinases, granzyme A-activated exonuclease, transcription factor, and GTPase regulation. The work presented here provides a detailed structural and energetic analysis of group I NDPKs, revealing the key position and contribution of the C-terminal sequence for dimeric interactions and hexameric stability. The superimposition of group I NDPK crystal structures highlights a high similarity between monomers and hexamers, albeit having divergent sequences and multiple oligomeric forms. We find the conserved Arginine residues to be critical for homohexameric assembly of group I NDPK subunits. Although single mutations to group I NDPKs (NME1/2: Arg27, NME3: Arg44, NME4: Arg60) alone do not disrupt the hexamer folding, they do destabilize complexes, as evident with reduced C_α_ RMSD. The NME1 and NME2 monomers have a single Arginine (Arg27), whereas NME3 and NME4 have multiple Arginine residues (NME3: Arg43, Arg44, Arg47; NME4: Arg60, Arg63, Arg64). We find that double and triple mutations in NME4 (containing shorter C-terminal regions) result in hexameric disassembly and propose that the increased stability of the NME1-3 hexamer is linked to the interaction between the C-terminal and *Kpn* loop of the adjacent monomer, lacking in NME4. For NME4, we observe that Arginine mutations lead to increased *Kpn* loop flexibility and an unstructured G91-D97 motif, which in turn perturbs the A108-V115 segment, disrupting interactions at the trimer interface. The identification of key arginine residues offers a new perspectives on oligomeric function, as mutation Arg27Gly has been shown to destabilize protein folding when overexpressed in *E.coli* [42]. In addition to single Arginine mutations, other factors like temperature and pH increase the susceptibility of the hexameric structure to disassembly. It is important to note that current MD trajectories are restricted to short durations with known hexameric structures, which may obscure the effects of intermolecular potential energy when performed over long simulation times. The combination of a truncated C-terminal region (Δ10ct) with Arginine mutations to NME1-3 (mimicking NME4) disrupts leads to a drastic reduction in binding free energy and hexamer disassembly (Table 1). Our findings are consistent with work from Kim et al., who reported strong interactions at the C-terminal of neighbouring monomers despite performing mutagenesis on the C-terminal region [31]. Similar observations had been shown in *Leishmania major,* where double mutation of Pro97Ser and shortened 5 residues at the C-terminal (Δ5Ct) affect the hexameric stability [53]. The reported human NME4 structure also shares features with *Leishmania braziliensis* NDK, which include a lack of interactions between the unstructured C-terminal and trimer interfaces, formed by Pro12, Asp13, Gln16, and Arg17. Furthermore, residues Val109-Arg113 are likely to contribute to the energetic stability of hexamers [54]. The NME1-3 C-terminal is ∼10 residues longer, and our work proposes that mutations (Δ10Ct) make the complex more labile and susceptible to denaturation (Fig. S8D). The contribution of C-terminal sequence to protein has been suggested in *D. melanogaster* [55], *Dictyostelium* [26, 56]. Notably in *M. tuberculosis*, the NME homology protein lacking C-terminal sequences exists as a thermostable hexamer (T_m_ of 76°C) with strong ionic interactions between neighbouring subunits due to entanglement with additional hydrophobic patches. Prokaryotic wildtype NME has also been reported to produce thermostable NDPK protein, where the side chain of the Cys133 forms disulfide bridges with neighbouring subunits [57]. The mutated NME4 homohexamer (double and trimer mutation) and its interface (monomerA to monomer D) stabilizes the dimer interface and, is prone to hexamer destabilization due to the lack of interaction between its C-terminal with the *Kpn* loop (NME1: 92-116) for its trimer interface. Crucially, our work also predicts NME1-2 heterohexamer configurations, which include a lower stability of NME1, NME2 homohexamer (A_6_B_0_, A_0_B_6_) compared to the penta-monomer combinations (A_5_B_1_, A_1_B_5_), where one monomer is swapped within the homohexamer assembly. The identification of heterohexamer configurations opens up exciting possibilities for purifying and understanding their contribution to a variety of NDPK roles. The possibility of novel post-translational modifications and interactions within heterohexamers offers valuable avenues for future research.

## Declarations

### Data Availability

The methods section contains the full data sources analysed in the current study. The analysis scripts detailing all the analysis steps can be found on GitHub repository (www.github.com/Natarajanlab/ndpk_oligo_states).

### Author contributions

KNN and YY designed the project. YY performed the analysis with inputs from KNN. KNN and YY wrote the manuscript. Both authors approve the manuscript.

## Acknowledgements

The research in the KNN lab is supported by the Villum Young Investigator grant (VYI#00025397) and DigitSTEM Initiative.

## Material and Methods

### Crystallized structure data collection and processing

We used available *Homo sapien* NME1-NME4 X-ray crystallography structures as starting point hexamer intermolecular interaction. For each group I NDPK, we extracted coordinates from Protein Data Bank (PDB) including NME1 (*1JXV*) [58], NME2 (*8PYW*) [59], NME3 (*1ZS6*) and NME4 (*1EHW*) [60], respectively.

Each crystal structure was corrected for polypeptide chain labelling for subsequent preparation of comparable quaternary structures. The different structure mutagenesis was performed using Chimera [61], and iteratively refined using the refinement package in AMBER [62]. For calculating surface electrostatic potential, we used APBS [63] on the trimer layer of different group I NDPKs.

### Computational alanine scanning mutagenesis (CASM)

The thermodynamic protein-protein binding affinity *ΔG_bind_* measures the strength of protein-protein interaction and is defined using the Gibbs free energy:

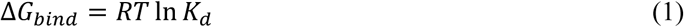

where, *R* represents the Boltzmann constant, *T* denotes the absolute temperature (in K), and the *K_d_* is the equilibrium dissociation constant of protein-protein interaction. By convention, a more negative *ΔG_bind_* value (expressed in kcal/mol) indicates a stronger interaction.

To determine the importance of residue’s positioning, we calculate changes in protein-protein binding affinity upon mutation (*ΔΔG_bind_*). This change in binding affinity due to a mutation is defined as:

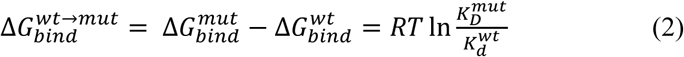

where, *wt* and *mut* refer to wildtype and mutant, respectively. Negative *ΔΔG_bind_* values correspond to mutations that stabilizes the complex upon mutation, while positive *ΔΔG_bind_* values indicate destabilization of the complex upon mutation.

### All-Atom Molecular Dynamics (MD) Simulations

For each homohexameric system, we apply a standard protocol using *tleap* module in Amber24 [62]. The Amber force field *ff14SB* [64] were assigned to all proteins and prior to solvation, either Na+ or Cl- counter ions were added to neutralize the net charge of each system. Each system is solvated in TIP3P water box [65], extending it with a minimum distance of 12 Å from the protein (solute) the surfaces of the octahedral water box in three dimensions. This solvation system and parameters mimic the natural cellular physiological environment. We performed all the all-atom MD simulations using AMBER24 package [62]. During the simulations, the system was first energy minimized by the steepest descent method for approximately 2000 cycles of steepest descent and 2000 cycles of conjugate gradient minimizations. We restrained protein backbone for a total of 4000 cycles of energy minimization with harmonic force constant of 50 kcal.mol^-1^.Å^-2^.

Then, the protein backbone is relaxed by turning off all the restraints then minimized for approximately 2000 cycles. Then the system was gradually heated from 0K (kelvin) to 310K in the in the NVT ensemble over a period of 1000 ps and then relaxed by 2000 ps in the NPT ensemble by Berendsen barostat with time step of 2fs. Finally, we perform 200 ns NPT simulations for each system without keeping any retraining spring constant. We employ the SHAKE algorithm [66] for all bonds involving hydrogen atoms. The particle mesh Ewald (PME) method was used when calculating the long-range electrostatic interactions [67], and the Lennard-Jones (LJ) potential. The van der Waals interactions were treated with twin range cut-off distance of 8Å and neighbour list cut-off of 10.0 Å. The periodic boundary conditions is imposed on all three directions (x, y, z).

### MM/PBSA and MM/GBSA calculations

A total of 20 frames with an interval of 10 ps in the final 200 ps of the MD trajectories were used for the calculation of total binding free energy (ΔG_bind_). All calculations were performed by using the *mmpbsa_py_energy* program to solve the Generalized-Born equations numerically. The ΔG_bind_ is calculated using the MM/GBSA methodology through the following equations:

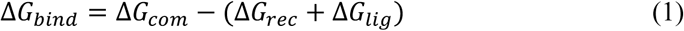

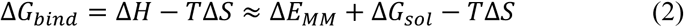

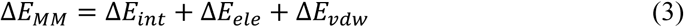

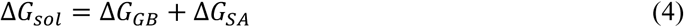

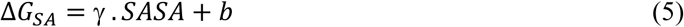

where ΔG_bind_ (total binding free energy) represents the free energy difference between ΔG_com_ (bound complex) and the sum of receptor (ΔG_rec_) and ligand (ΔG_lig_), respectively. In addition, it can be decomposed into three terms: gas-phase interaction energy (ΔE_MM_), desolvation energy (ΔG_sol_), and conformation entropy (-TΔS). The conformation entropy (-TΔS) however, was not calculated due to its high computational cost and low prediction accuracy [68]. ΔE_MM_ contains ΔE_int_ (bond, angle, and dihedral energies), ΔE_ele_ (electrostatic) and ΔE_vdw_ (van der Waals). Meanwhile, ΔG_sol_ is the sum of non-polar contribution (ΔG_SA_) estimated using the solvent accessible surface area (SASA) and polar contribution (ΔG_GB_) calculated by using the Generalized Born (GB) model. The solute interior dielectric constant (ε_int = 2_) was employed for the calculation. Meanwhile, the exterior (solvent) dielectric constant (χ_ext_) was set to 80. The non-polar component of desolvation energy was estimated by using the LCPO algorithm [69] and the formal surface area is usually estimated using the solvent accessible surface area (SASA), where ψ and 4 were set to 0.0072 kcal.mol^-1^.Å^-2^ and 0 kcal.mol^-1^, respectively.

